# Combining CRISPR/Cas9 mutagenesis and a small-molecule inhibitor to probe the function of MELK in cancer

**DOI:** 10.1101/203984

**Authors:** Christopher J. Giuliano, Ann Lin, Joan C. Smith, Ann C. Palladino, Jason M. Sheltzer

**Author notes:** Denotes equal contribution (Christopher J. Giuliano, Ann Lin).

## Abstract

The Maternal Embryonic Leucine Zipper Kinase (MELK) has been identified as a promising therapeutic target in multiple cancer types. MELK over-expression is associated with aggressive disease, and MELK has been implicated in numerous cancer-related processes, including chemotherapy resistance, stem cell renewal, and tumor growth. On the basis of these findings, a MELK inhibitor is currently being tested in several clinical trials. Here, we report that cancer cell lines harboring CRISPR/Cas9-induced null mutations in MELK exhibit wild-type growth *in vitro*, under environmental stress, in the presence of multiple chemotherapy agents, and *in vivo*. By combining our MELK-knockout clones with a recently-described, highly-specific MELK inhibitor, we further demonstrate that the acute inhibition of MELK results in no specific anti-proliferative phenotype. Analysis of gene expression data from cohorts of cancer patients identifies MELK expression as a correlate of tumor mitotic activity, explaining its association with poor clinical prognosis. In total, our results demonstrate the power of CRISPR/Cas9-based genetic approaches to investigate cancer drug targets, and call into question the rationale for treating patients with anti-MELK monotherapies.

## Introduction

Cancer cells require the expression of certain genes, called “addictions” or “genetic dependencies”, that encode proteins necessary for tumor growth. Silencing the expression of these genes or blocking the activity of the proteins that they encode can trigger cell death and durable tumor regression (1). Identifying and characterizing cancer dependencies is therefore a key goal of pre-clinical cancer research.

The Maternal Embryonic Leucine Zipper Kinase (MELK) has been implicated as a cancer dependency and putative drug target in multiple cancer types, including melanoma, colorectal cancer, and triple-negative breast cancer (2–6). MELK is over-expressed in these cancers, and high expression of MELK is associated with poor patient prognosis (4, 7–10). Moreover, knockdown of MELK using RNA interference (RNAi) has been reported to block cancer cell proliferation and trigger cell cycle arrest or mitotic catastrophe (3, 4, 10–13). On the basis of these pre-clinical results, several companies have developed small-molecule MELK inhibitors, and one MELK inhibitor (OTS167) is currently being tested in multiple clinical trials (14).

In contrast to these results, we recently reported that triple-negative breast cancer cells harboring CRISPR/Cas9-induced loss of function mutations in MELK proliferate at wild-type levels *in vitro* (15). Additionally, we demonstrated that the MELK inhibitor OTS167 remained effective against MELK-mutant cells, suggesting that OTS167 kills cells through an off-target mechanism. These results have been replicated by an independent group, who further demonstrated that the shRNA vectors commonly used to study MELK also kill cells in a MELK-independent manner (16). The off-target effects of both the small-molecule MELK inhibitor and the MELK-targeting shRNAs provide a potential explanation for certain previous results obtained studying this potential drug target.

Despite the conflicting *in vitro* data, MELK expression remains one of the strongest predictors of patient mortality in diverse cancer types (17). Additionally, MELK has been implicated in several other cancer-related processes, including cancer stem cell maintenance, chemotherapy resistance, anchorage-independent growth, and reactive oxygen species (ROS)-signaling (4, 5, 13, 18–22). These processes may not be challenged by the routine *in vitro* growth assays that have been performed in MELK-knockout (MELK-KO) cells to date. Several of these studies were conducted using distinct RNAi constructs and small-molecule inhibitors, raising the possibility that they reflect true functions of this kinase. Moreover, the over-expression of MELK has been reported to transform cells, suggesting that in addition to MELK’s putative role as a cancer dependency, it may also function as a driver oncogene (4).

While RNA interference is susceptible to off-target interactions that confound experimental interpretation (23, 24), CRISPR/Cas9 mutagenesis is also prone to several important limitations. In particular, clonal cell lines harboring Cas9-induced modifications must be expanded from a single cell with a mutation of interest to several million cells. This intense pressure may select for secondary mutations that blunt any anti-proliferative consequences of the original mutation (25). In a therapeutic context, the immediate inhibition of a particular target achieved with a small-molecule drug may induce a more severe phenotype than observed in a CRISPR-modified cell line subjected to evolutionary pressure over the course of days or weeks.

To investigate the role of MELK in cancer-related processes beyond cell proliferation, and to assess the therapeutic potential of immediate MELK inhibition, we performed assays combining CRISPR-knockout cell lines with a recently-described, highly-specific MELK inhibitor (16). In a variety of *in vitro* and *in vivo* challenges, we found that cells lacking MELK behave indistinguishably from wild-type cells. Moreover, through a close analysis of gene expression data, we report that MELK levels strongly correlate with mitotic activity in human tumors, suggesting that MELK may function as an indirect proxy for aggressive growth. In total, these results cast doubt on the possibility that MELK-specific inhibition will serve as a useful monotherapy in cancer.

## Results

### MELK over-expression fails to transform immortalized cell lines

The over-expression of driver oncogenes allows immortalized but non-transformed cell lines to form colonies when grown in soft agar, a phenotype that is tightly linked with *in vivo* tumorigenicity (26–28). It has previously been reported that the over-expression of MELK was sufficient to induce anchorage-independent growth in several cell lines, including Rat1 fibroblasts expressing dominant-negative p53 (p53dd) and the human mammary epithelial cell line MCF10A (4). We attempted to replicate these results using Rat1-p53dd and MCF10A cells, as well as the immortalized 3T3 mouse fibroblast cell line. To accomplish this, we stably transduced each cell line with a retroviral vector encoding either the mouse or the human MELK protein. Western blot analysis confirmed that full-length mouse or human MELK was over-expressed in all six cell lines that we generated relative to the level of MELK in vector-transduced control cell lines (Figure 1A,D). Additionally, as positive controls in this assay, we transduced each cell line with an allele of H-Ras known to function as a strong driver oncogene (H-Ras^G12V^)(29), and with an allele of EGFR that weakly transforms cells (EGFR^L858R^) (30). We then assessed whether cell lines over-expressing each gene would proliferate when suspended in soft agar. As expected, cells that had been transduced with an empty vector exhibited minimal anchorage-independent growth, while cells transduced with H-Ras^G12V^ or EGFR^L858R^ formed numerous colonies (Figure 1B-C and E-F). However, in multiple independent experiments, we failed to detect an increase in anchorage-independent growth in any of the six cell lines over-expressing either mouse or human MELK. We conclude that, under the conditions tested, the over-expression of MELK fails to transform cells.

**Figure 1.**
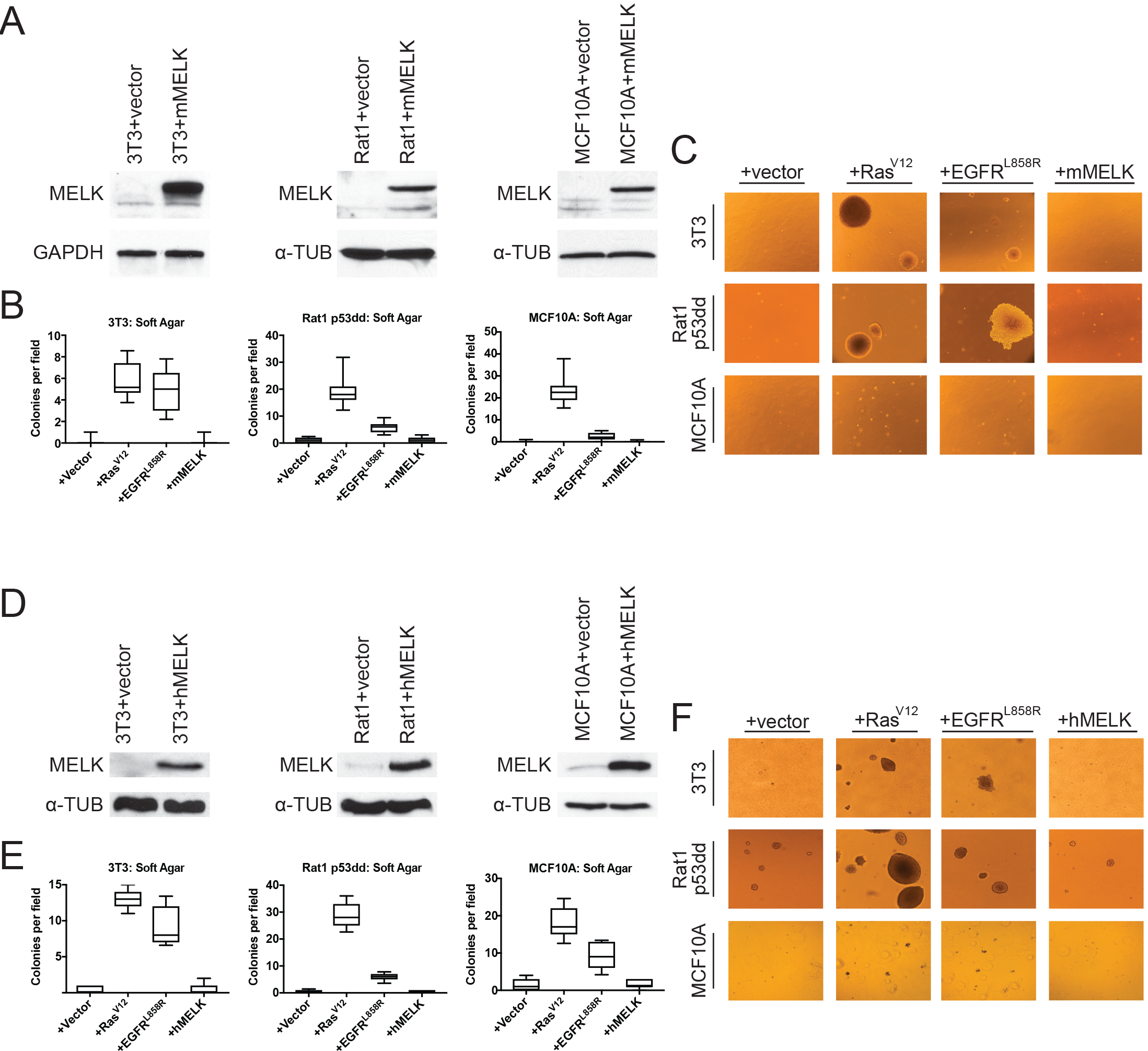
MELK over-expression fails to confer anchorage-independent growth. (A) Western blot analysis of mouse MELK over-expression in 3T3, Rat1-p53dd, and MCF10a cell lines. (B) Quantification of colony formation of control and mouse MELK over-expressing cell lines in soft agar. For each assay, colonies were counted in at least 15 fields under a 10× objective. Boxes represent the 25th, 50th, and 75th percentiles of colonies per field, while the whiskers represent the 10th and 90th percentiles. (C) Representative images of the indicated cell lines grown in soft agar. (D) Western blot analysis of human MELK over-expression in 3T3, Rat1-p53dd, and MCF10 cell lines. (E) Quantification of colony formation of control and human MELK over-expressing cell lines in soft agar. For each assay, colonies were counted in at least 15 fields under a 10× objective. Boxes represent the 25th, 50th, and 75th percentiles of colonies per field, while the whiskers represent the 10th and 90th percentiles. (F) Representative images of the indicated cell lines grown in soft agar.

### MELK is dispensable for growth *in vitro* and *in vivo*

We previously showed that MELK was not required for cell division in two triple negative breast cancer cell lines, Cal51 and MDA-MB-231 (15). However, MELK has been reported to support growth in several other cancer types, including colorectal cancer and melanoma (6, 10, 20). To examine the role of MELK in other cancer types, we used CRISPR/Cas9 to generate multiple MELK-knockout (MELK-KO) clones in A375, a melanoma cell line, and DLD1, a colorectal cancer cell line (Supplemental Figure 1). As controls, we also derived clones of A375 and DLD1 harboring guide RNAs that targeted the non-essential Rosa26 locus. MELK mutagenesis was verified by sequencing the sites targeted by the gRNA, and loss of the MELK protein was verified by western blot (Supplemental Figure 1A-C). MELK-KO melanoma and colorectal cancer clones grew at wild-type levels *in vitro*, demonstrating that MELK is dispensable for proliferation in these cancer types as well (Supplemental Figure 1D-E). We previously reported that OTS167, a putative MELK inhibitor in clinical trials, kills breast cancer cells in a MELK-independent manner (15). Consistent with these observations, MELK-KO and Rosa26 clones were equally sensitive to OTS167, verifying that this drug kills cells via an off-target effect across cancer types (Supplemental Figure 1F).

Many genes that are non-essential for cell division *in vitro* may still play crucial roles in cancer by supporting other processes, including stem-cell renewal, resistance to anoikis, and angiogenesis (31–34). We therefore subjected our MELK-KO and Rosa26 clones to various *in vitro* and *in vivo* assays to assess whether MELK loss impairs any cancer-related phenotypes. Although all MELK-KO cells grow well when seeded at high density in proliferation assays (15, 16), plating cells at low density can challenge a cell’s colony-forming ability and replicative lifespan (35). To test whether MELK loss confers a defect in colony growth, MELK-KO and Rosa26 DLD1, A375, Cal51, and MDA-MB-231 clones were serially-diluted and allowed to grow at varying cell densities. Crystal violet staining of these plates revealed that the loss of MELK failed to impair colony growth relative to the control Rosa26 cell lines (Figure 2A). In fact, one MELK-KO clone (MDA-MB-231 c1) grew consistently better than either control clone in this assay. Significant variation in proliferative capacity has previously been described among independent clones of the MDA-MB-231 cell line (36), and may arise due to heterogeneity in the parental population or secondary mutations acquired during cell line derivation.

**Figure 2.**
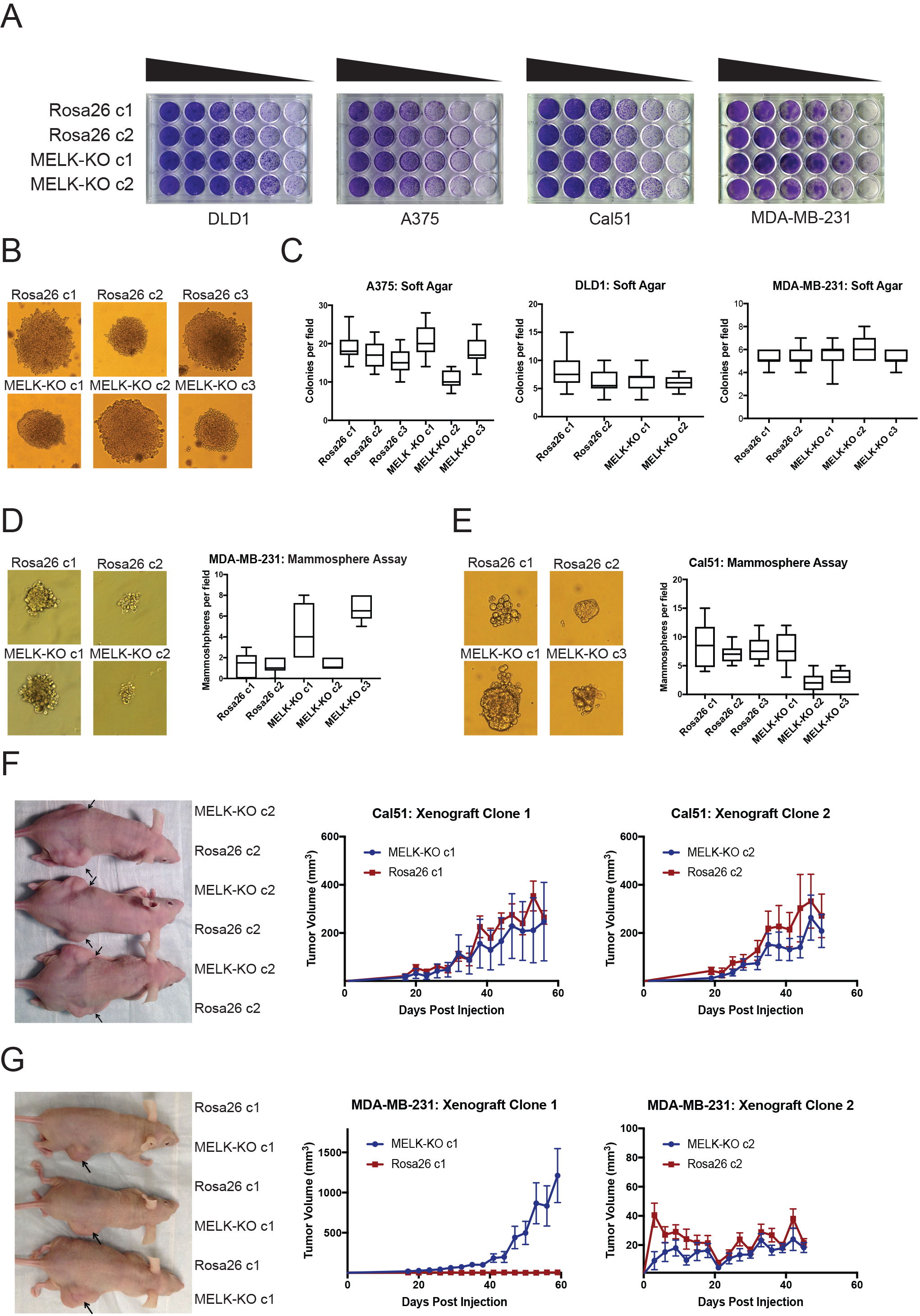
MELK is dispensable for growth *in vitro* and *in vivo*. (A) Crystal violet staining of serial dilution plates of control and MELK knockout clones from DLD1 (colorectal cancer), A375 (melanoma), Cal51 (breast cancer) and MDA-MB-231 (breast cancer) cell lines. (B) Representative images of colonies of A375 control and MELK knockout clones grown in soft agar. (C)Quantification of colony formation in A375, DLD1, and MDA-MB-231 control and MELK knockout clones. For each assay, colonies were counted in at least 15 fields under a 10× objective. Boxes represent the 25th, 50th, and 75th percentiles of colonies per field, while the whiskers represent the 10th and 90th percentiles. (D-E) Representative images and quantification of mammosphere growth in MDA-MB-231 or Cal51 control and MELK knockout clones. For each assay, mammospheres were counted in at least 6 fields under a 10x objective. Boxes represent the 25th, 50th, and 75th percentiles of colonies per field, while the whiskers represent the 10th and 90th percentiles. (F-G) Representative images and quantification of xenograft growth in nude mice. Cal51 and MDA-MB-231 control and MELK knockout clones were injected subcutaneously into nude mice, and then tumor growth was measured every three days. Arrows indicate the location of the tumor. Error bars in the volume measurements indicate the standard error.

To further investigate the impact of MELK loss on anoikis and tumorigenicity, MELK-KO and Rosa26 cells were plated in soft agar. However, we observed no significant difference in anchorage-independent growth between MELK-KO cells and Rosa26 cells in every cell line tested (Figure 2B-C). MELK has previously been implicated in the maintenance of breast cancer stem cells (5, 21, 37). We therefore tested whether loss of MELK impairs mammosphere formation, a phenotype tightly linked with breast cancer stem cell activity (38). While we observed inter-clonal variability in mammosphere growth, all MELK-KO clones were capable of forming mammospheres, and two MELK-KO clones exhibited consistently greater mammosphere formation than their wild-type controls (Figure 2D-E). We conclude that MELK is dispensable for growth as a mammosphere.

Finally, we sought to determine whether MELK was required for tumor formation *in vivo*. To address this, we performed flank injections into nude mice with multiple clones of Cal51 and MDA-MB-231 MELK-KO and Rosa26 cells. Across all clones tested, 23 of 34 injections with MELK-KO cells and 24 of 34 injections with Rosa26 cells resulted in detectable tumor formation, and no significant growth defect was observed in any MELK-KO clone (Figure 2F-G). Consistent with our previous assays, we observed superior growth in the MDA-MB-231 MELK-KO c1 clone, while MDA-MB-231 MELK-KO c2 and Rosa26 c2 grew at equivalent rates (Figure 2G). In total, our results demonstrate that MELK is dispensable for the proliferation of cancer cells *in vitro* and *in vivo*.

### MELK is not required for the phosphorylation or expression of previously reported targets

MELK has been reported to support cancer cell proliferation by phosphorylating various proteins involved in splicing, translation, metabolism, and cell cycle progression (19, 39–42). In particular, a recent publication reported that MELK phosphorylates eukaryotic translation initiation factor 4B (eIF4B), and this phosphorylation event promotes cell survival by increasing translation of the anti-apoptotic protein MCL1 (39). However, western blot analysis revealed normal levels of eIF4B phosphorylation in MELK-knockout MDA-MB-231, CAL51, and A375 cells (Supplemental Figure 2A-C). Additionally, MELK-KO cell lines continued to express MCL1, the putative downstream target of eIF4B (Supplemental Figure 2 D-E). We conclude that MELK is not required for eIF4B phosphorylation or MCL1 translation.

### MELK is dispensable for cell growth under exogenous stress

Developing tumors must survive in hypoxic and nutrient-poor conditions (43), and MELK has been implicated in glucose signaling and in the detection of ROS (19, 44). We therefore considered the possibility that MELK expression is necessary to support growth under metabolic or environmental stress. To generate ROS stress, we cultured cells in varying concentrations of H_2_O_2_, but we observed no difference in sensitivity between MELK-KO and control Rosa26 clones (Figure 3A). MELK has been suggested to contribute to ROS signaling by phosphorylating ASK1 (44); however, this protein remained phosphorylated at normal levels in cells lacking MELK (Supplemental Figure 2F). MELK-KO clones also exhibited wild-type levels of growth when cultured under hypoxic, serum-deprived, or glucose-limited conditions (Figure 3B-D). We conclude that MELK is dispensable for proliferation under common metabolic stresses.

**Figure 3.**
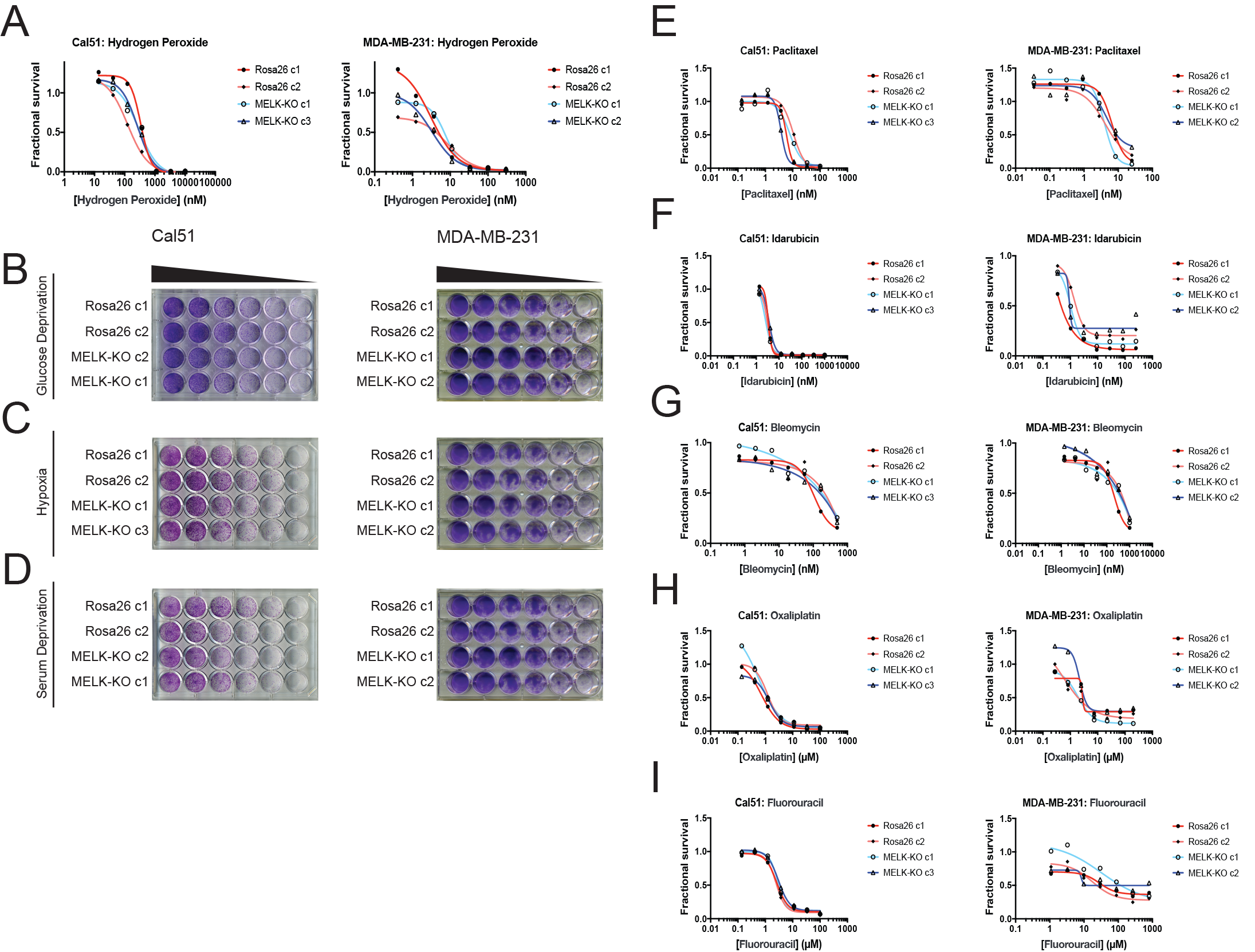
MELK is not required for growth under stress. (A) Dose-response curves of Cal51 and MDA-MB-231 Rosa26 and MELK-KO clones grown in the presence of H_2_O_2_. (B-D) Crystal violet staining of Cal51 and MDA-MB-231 Rosa26 and MELK-KO clones grown as serial dilutions under the indicated stressful culture condition. (E-I) Dose-response curves of Cal51 and MDA-MB-231 Rosa26 and MELK-KO clones grown in the presence of the indicated chemotherapy drug.

As high MELK expression is associated with poor patient prognosis, we wondered if MELK could promote resistance to cytotoxic chemotherapies. Indeed, it has been previously reported that knocking down or inhibiting MELK sensitizes cells to DNA damage (13, 18, 20, 37). To test whether MELK has a role in chemotherapy resistance, we performed drug sensitivity assays using a variety of DNA-damaging or anti-mitotic agents. However, we observed no significant difference in viability between MELK-KO and Rosa26 cell lines treated with five common chemotherapies (Figure 3E-I). In total, these results demonstrate that MELK is dispensable for cell survival under metabolic and cytotoxic stress.

### Acute inhibition of MELK fails to block proliferation

Deriving CRISPR-knockout clones from single cells selects for a population of cells that are capable of surviving clonal expansion. Since our MELK-KO cell lines were generated from single cells, we considered the possibility that MELK plays an important role supporting proliferation, but the clones we generated had evolved to tolerate the loss of MELK. To assess this possibility, we performed an epistasis experiment combining our MELK-knockout clones with a recently-described, highly-specific small molecule MELK inhibitor, HTH-01-091 (16). We reasoned that treating Rosa26 clones with HTH-01-091 would reveal the consequences of the acute loss of MELK. However, if such phenotype(s) were also present in MELK-KO cell lines treated with HTH-01-091, then the phenotype(s) could be attributed to an off-target effect of the drug.

We first sought to identify a concentration at which HTH-01-091 inhibited MELK in our cells of interest. However, as described in this manuscript, we lacked a verified MELK substrate whose phosphorylation status could be monitored to confirm MELK inhibition. Nonetheless, it has been reported that a by-product of MELK inhibition is the degradation of MELK protein (13, 16). Therefore, to determine an effective concentration of HTH-01-091, we monitored the level of MELK protein after drug treatment by western blot. We found that 1μM HTH-01-091 triggered near-complete MELK degradation in MDA-MB-231, DLD1, and Cal51 cells (Figure 4A). The loss of MELK was not an indirect effect of cell cycle arrest, as these cells maintained high levels of the mitotic marker cyclin B. This concentration was therefore used in subsequent assays.

**Figure 4.**
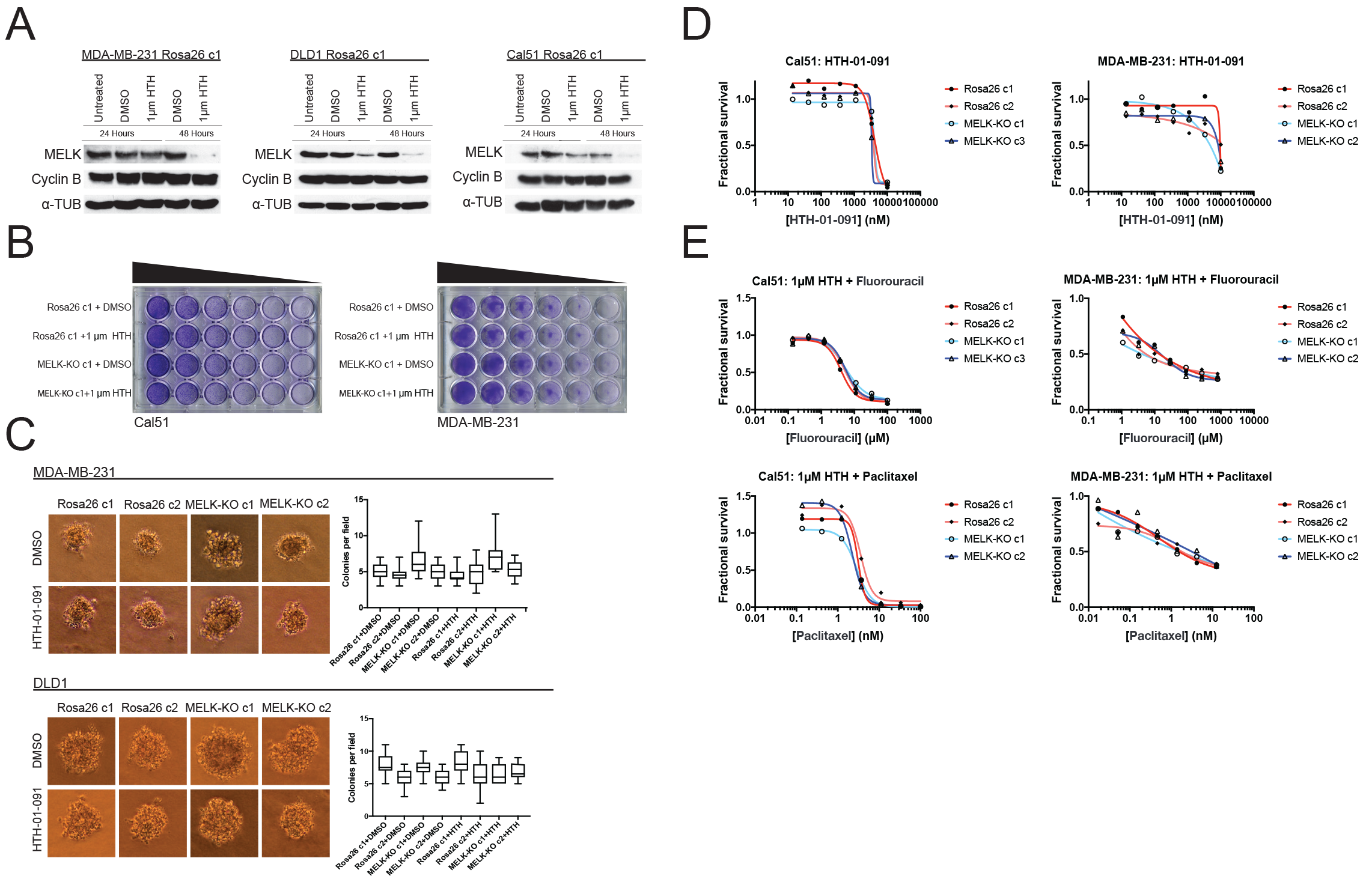
Acute inhibition of MELK fails to block growth. (A) Western blot analysis of MELK expression levels during treatment with 1μM HTH-01-091 in the MDA-MB-231, DLD1, and Cal51 Rosa26 clonal cell lines. (B) Crystal violet staining of Cal51 and MDA-MB-231 Rosa26 and MELK-KO cell lines grown as serial dilutions in the presence of DMSO or 1μM HTH-01-091. (C) Quantification and representative images of colony formation of MDA-MB-231 and DLD1 Rosa26 and MELK-KO clones in soft agar in the presence of DMSO or 1μM HTH-01-091. (D)Dose-response curves of Cal51 and MDA-MB-231 Rosa26 and MELK-KO clones grown in the presence of HTH-01-091. (E) Dose-response curves of Cal51 and MDA-MB-231 Rosa26 and MELK-KO clones grown in the presence of 1μm HTH-01-091 and the indicated chemotherapy drug.

To test whether the acute inhibition of MELK affected clonogenicity or anchorage-independent growth, we grew MELK-KO and Rosa26 cells on plastic or in soft agar in the presence of DMSO or 1μM HTH-01-091. Neither MELK-KO nor Rosa26 clones were affected byHTH-01-091 treatment, verifying that MELK is dispensable for colony formation and anchorage-independent growth in these cells (Figure 4B-C). Indeed, a drug sensitivity assay revealed that HTH-01-091 exhibited significant anti-proliferative effects only at concentrations above ~5μM (Figure 4D). This toxicity is likely a consequence of an off-target effect, as these drug concentrations were found to affect Rosa26 and MELK-KO cells equivalently.

We next sought to test whether acute MELK inhibition sensitized cells to chemotherapy. To accomplish this, we treated MELK-KO and Rosa26 clones with various chemotherapy drugs in the presence of HTH-01-091. Consistent with our previous results, HTH-01-091 treatment failed to sensitize the Rosa26 clones to 5-florouracil or paclitaxel treatment (Figure 4E). Finally, we assessed the effect of HTH-01-091 treatment on eIF4B phosphorylation and MCL1 expression, and found that MELK inhibition failed to affect either target (Supplemental Figure 2G-H). In total, these results demonstrate that the acute loss of MELK results in no significant defect in cell viability, proliferation, or drug resistance, and suggest that our knockout clones have not acquired mutations that tolerize cells to the loss of MELK.

### The association between MELK expression and cancer lethality is due to its correlation with mitotic activity

Our *in vitro* and *in vivo* experiments failed to reveal any cancer-related phenotypes affected by either the deletion or over-expression of MELK. Yet, MELK is up-regulated in many cancer types (10), and high levels of MELK expression have been reported to confer a dismal clinical prognosis (4, 7–9). If MELK plays no overt role in cancer biology, then why would MELK expression be linked with death from cancer? We note that MELK expression is cell cycle-regulated, peaking in mitosis (4, 45), and gene signatures that capture mitotic activity have been found to be prognostic in multiple cancer types (17, 46, 47). We therefore considered the possibility that, rather than functioning as an oncogene or a cancer dependency, MELK expression could report cell division within a tumor. To assess the link between MELK expression and cell proliferation, we analyzed gene expression data from different sets of cells and tissues. In normal human tissue, MELK expression was the lowest in non-proliferative organs, including the heart and skeletal muscle, while MELK expression was the highest in organs with on-going mitotic activity, including the bone marrow and testes (Figure 5A). Indeed, across 32 tissue types, MELK levels were highly correlated with the expression of the proliferation marker MKI67 (R = 0.93). Similarly, in human fibroblasts cultured until senescence, MELK expression decreased up to 11-fold between proliferating and arrested populations, while stimulating lymphocytes to divide increased MELK expression 20-fold (Figure 5B-C). These data suggest that MELK levels reflect mitotic activity in diverse cell types.

**Figure 5.**
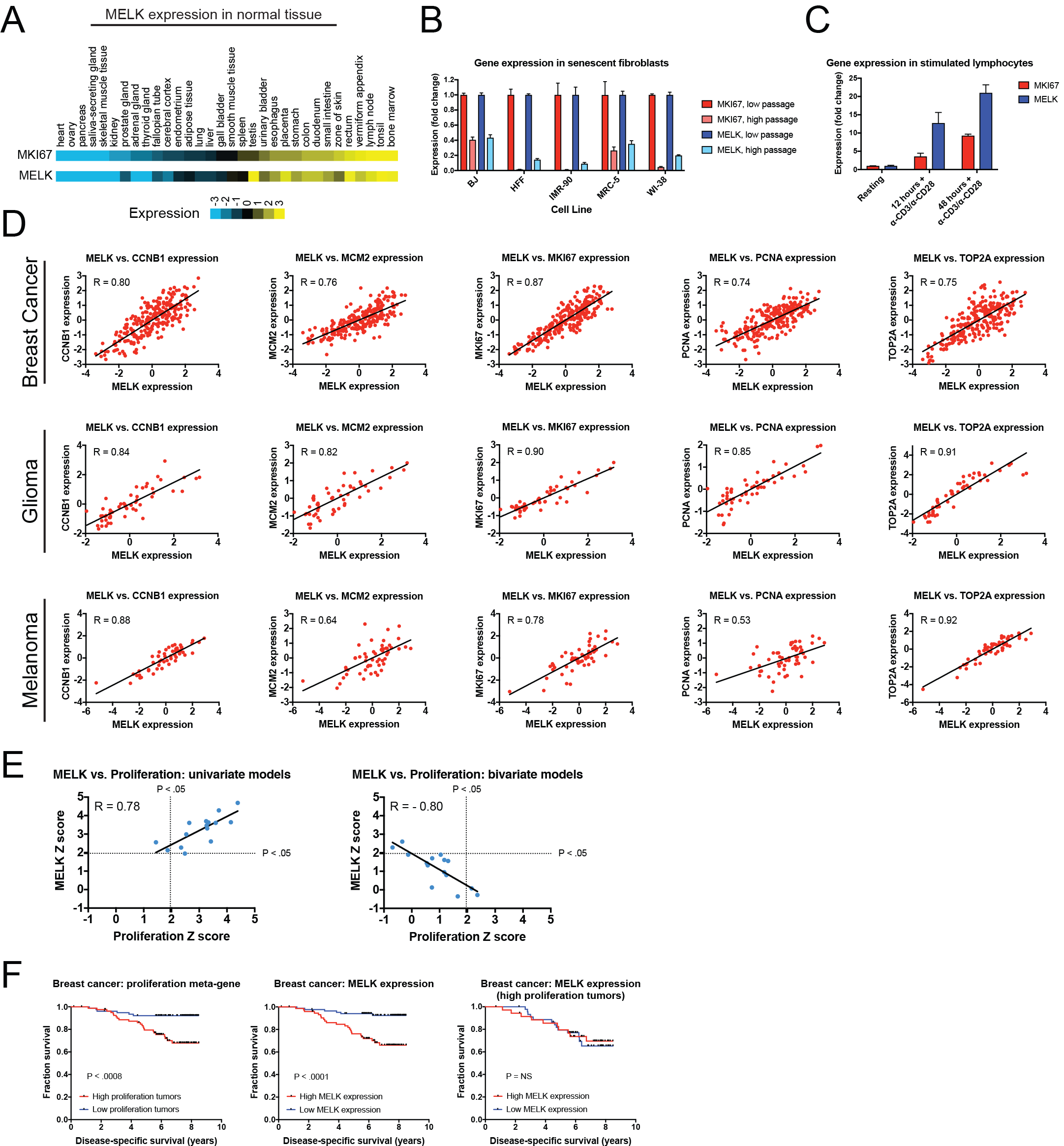
MELK expression correlates with proliferation markers *in vitro*, in normal tissue, and in cancer. (A) A heatmap of the expression of either MKI67 or MELK in normal tissue sorted according to MKI67 expression (53). (B) The expression level of either MKI67 or MELK is displayed in five different primary human fibroblast lines at either low passage (proliferating) or high passage (senescence) (54). (C) The expression level of either MKI67 or MELK is displayed in CD4+ lymphocytes resting or after stimulation with α-CD3/α-CD28 beads (55). (D) The expression level of MELK is plotted against the expression of five common proliferation markers in cohorts of patients with breast cancer, glioma, or melanoma (57–59). Black lines represent linear regressions plotted against the data. (E) Univariate (left) or bivariate (right) Cox proportional hazards models were calculated for the 15 breast cancer cohorts listed in Table S3 and S4. For each cohort, the expression of either MELK or the average expression of CCNB1, MCM2, MKI67, PCNA, and TOP2A in each tumor was regressed against patient outcome. Dotted lines represent Z scores of 1.96, corresponding to a p-value of 0.05. Black lines represent linear regressions plotted against the data. (F) Kaplan-Meier curves displaying disease-specific survival in one breast cancer cohort (60). Patients were split into two populations based on the average expression of either MELK or the five-gene proliferation meta-gene.

To examine the link between MELK and cell division in cancer, we compared the levels of MELK expression with 5 well-characterized proliferation markers: MKI67, PCNA, CCNB1, MCM2, and TOP2A (48). In cohorts of patients with tumor types in which MELK levels are associated with advanced disease, MELK expression was significantly correlated with each of the proliferation genes (median correlation = 0.82; Figure 5D). We then sought to determine whether the correlation between MELK expression and proliferation could explain the prognostic significance of MELK. To test this, we collected 15 breast cancer microarray datasets from patients with known clinical outcomes. We calculated Z scores for univariate Cox models for each patient cohort, and found that MELK expression was associated with poor outcome in 14 of 15 datasets (Table S3). Similarly, a proliferation meta-gene derived by combining MKI67, PCNA, CCNB1, MCM2, and TOP2A was prognostic in 13 of 15 datasets (Table S3). Strikingly, we found that the Z scores generated by our univariate proliferation model were highly correlated with the Z scores generated by the univariate MELK model (Figure 5E; R = 0.78, P < .001), suggesting that these two models capture very similar clinical information. To confirm this, we constructed bivariate Cox models that considered both MELK expression and the proliferation meta-gene (Table S4). In the bivariate models, MELK was significantly associated with patient outcome in only 2 of 15 datasets, demonstrating that considering tumor cell proliferation ablated its clinical utility (Figure 5E). Thus, when tumors are stratified according to their proliferation level, MELK expression is no longer prognostic (Figure 5F). In total, these results suggest that the observed pattern of MELK expression in cancer can be explained by the fact that MELK is up-regulated in rapidly-dividing cells.

## Discussion

Pre-clinical cancer research efforts apply different genetic and chemical tools (RNA interference, CRISPR, and small-molecule inhibitors) to a variety of artificial assays (*in vitro* proliferation, xenograft growth, etc.) in order to discover targets that will have clinical efficacy when inhibited in human patients. No single assay perfectly mimics the behavior of a tumor in a human cancer patient, and no single chemical or genetic tool exhibits absolute specificity. Nonetheless, we believe that by using multiple orthogonal approaches and assays, we can gain insight into the role that certain genes play in human malignancies. In this current manuscript, we report that combining CRISPR and a small-molecule inhibitor in a variety of assays failed to reveal a role for MELK in several cancer-related processes. These results suggest that anti-MELK monotherapies are unlikely to be effective cancer treatments.

Our previous work demonstrated that MELK is dispensable for the proliferation of triplenegative breast cancer cells *in vitro* (15). Nonetheless, multiple cellular functions are dispensable for *in vitro* proliferation but required for tumor progression, including stem cell renewal, oxygen sensing, and chemotherapy resistance (31–34). Though MELK has been implicated in each of these processes, our results demonstrate that MELK-knockout cancer cell lines grow at wild-type levels in a variety of assays designed to test these pathways. We speculate that, as has previously been reported for one MELK inhibitor and one set of MELK-targeting shRNA’s (16), several previous studies of MELK function may have been compromised by off-target activity of the constructs and inhibitors that were used.

To assess MELK function in cancer, we combined clonal cell lines harboring null mutations in MELK with a specific MELK inhibitor. Of note, we observed some variability between clonal lines, underscoring the importance of assessing multiple independent clones in CRISPR experiments. To rule out the possibility that our MELK-KO clones had evolved to tolerate the loss of MELK, we performed “epistasis” experiments by treating these knockout clones with a MELK inhibitor. We reasoned that specific consequences of MELK inhibition would be detectable upon drug treatment in MELK-WT but not MELK-KO clones, while nonspecific consequences of drug treatment would affect both genotypes equally. In all experiments conducted thus far, no phenotypes have been observed only in MELK-WT cells after HTH-01-091 treatment, further verifying that MELK is dispensable for cancer cell growth. We suggest that these “epistasis” experiments can be widely applied to assess the on-target consequences of acutely inhibiting potential cancer drug targets.

Initial interest in blocking MELK function in cancer stemmed from the discovery that it was over-expressed across cancer types (10). Further research revealed that patients whose tumors expressed the highest levels of MELK had the worst clinical outcomes (4, 7–9, 11). As we observed no role for MELK as either an oncogene or a cancer dependency, we sought to instead investigate whether its cell cycle-dependent expression pattern could explain its prognostic value. Consistent with this hypothesis, we discovered that MELK expression closely mirrors cell and tumor mitotic activity. As actively-growing tumors appear to up-regulate thousands of genes involved in cell cycle progression when compared to quiescent normal tissue (48), this observation may explain why MELK is commonly over-expressed in cancer. Moreover, rapid cell division in tumors is indicative of tissue de-differentiation and aggressive disease; therefore, most cell cycle-regulated genes are also associated with clinical outcome (17, 46, 47). We found that controlling for cell proliferation ablates the significance of MELK expression, suggesting that this link may explain its prognostic role in cancer.

While many cell cycle genes are indeed suitable cancer drug targets (e.g., CDK4 and CDK6), other genes may be up-regulated during normal cell division but dispensable for this process (46). We suggest that MELK is an example of the latter. We believe that, if MELK does play a role in cancer, it may be detectable only in very limited circumstances, and likely *in vivo*. As MELK-knockout mice display no observable deficiencies, MELK’s function may be redundant with other kinases (4). Future synthetic lethal screening and additional *in vivo* assays may clarify what role, if any, MELK plays in cancer biology.

## Acknowledgments

We thank Hubert Huang and Nathanael Gray (Dana-Farber Institute) for providing HTH-01-091. This work was performed with assistance from CSHL Shared Resources, including the CSHL Flow Cytometry Shared Resource, which are supported by the Cancer Center Support Grant 5P30CA045508. Research in the Sheltzer Lab is supported by an NIH Early Independence Award (1DP5OD021385), a Breast Cancer Alliance Young Investigator Award, and a CSHL-Northwell Translational Cancer Research Grant.

## Materials and Methods

### Cell lines and culture conditions

The identity of each human cell line was verified by STR profiling (University of Arizona Genetics Core). MDA-MB-231, Cal51, A375, and DLD1 cell lines were grown in DMEM supplemented with 10% FBS, 2mM glutamine, and 100 U/mL penicillin and streptomycin. Rat1 cells were grown in DMEM supplemented with 5% FBS, 2mM glutamine, and 100 U/mL penicillin and streptomycin. 3T3 cells were grown in DMEM supplemented with 10% bovine calf serum (BCS), 2mM glutamine, and 100 U/mL penicillin and streptomycin. MCF10A cells were grown in Mammary Epithelial Cell Growth Medium (Lonza; Cat. No. CC-3150) supplemented with 5% horse serum, 100 ng/mL of cholera toxin (Sigma-Aldrich; Cat. No. C8052) and 100 U/mL penicillin and streptomycin. All cell lines were grown in a humidified environment at 37°C and 5% CO_2_

### Retroviral plasmid over-expression

Mouse and human MELK cDNA was cloned into an MMLV vector and verified by sequencing (Vectorbuilder, Cyagen Corporation). Positive and negative control plasmids were acquired from Addgene: pBabe-Puro (Addgene; Cat. No. 1764), EGFR L858R (Addgene; Cat. No. 11012), and RasV12 (Addgene; Cat. No. 1768). Retrovirus was generated by transfecting plasmids into Plat-A cells (Cell BioLabs; Cat. No. RV-102) using the calcium-phosphate method (49). Virus was harvested 48-72 hours post transfection, filtered through a 0.45μm syringe and applied to cells with 4μg/mL polybrene. After 24 hours, the media was changed and cells were allowed to recover in fresh media for 2 days. Subsequently, the cells were split and the appropriate antibiotic was added to select for transduced cells.

### Soft agar assays

To assay anchorage-independent growth, all cell lines except MCF10A were suspended at a cell count of 10,000 cells in a 0.35% Difco Agar Noble (VWR Scientific; Cat. No. 90000-772) solution in a six-well plate. MCF10A cell lines were suspended at a cell count of 20,000 cells. The mixture was plated over a 0.5% Difco Agar Noble solution. Plates were allowed to solidify at room temperature for 2 hours and then placed in a 37°C incubator overnight. 1 mL of normal growth media was added to each well the next day and every 3 days after (50). After 14 days, colony formation was quantified under 20× magnification.

### Stress assays

To study the role of MELK in surviving stressful culture conditions, 30,000 MELK-KO and Rosa26 cells from MDA-MB-231 and DLD1 cell lines and 10,000 MELK-KO and Rosa26 cells from Cal51 and A375 cell lines were plated in the first column of a 24-well plate (Corning; Cat. No. 3526) and then five three-fold dilution were performed across the plate. For the low glucose conditions, low glucose DMEM (Thermo Fisher Scientific; Cat. No. 11885076) supplemented with 10% FBS, 2 mM glutamine, and 100 U/mL penicillin and streptomycin was added to the cells. For the low serum conditions, cells were cultured in DMEM supplemented with 1% FBS, 2 mM glutamine, and 100 U/mL penicillin and streptomycin. For the hypoxic conditions, cells were cultured in normal media and then placed in a hypoxic incubator set at 37°C with 2% oxygen. After 10-14 days, cells were fixed with 100% methanol and stained with 0.5% crystal violet dissolved in 25% methanol.

### Mammosphere formation assay

Mammosphere formation media was prepared using DMEM/F12 (Lonza; Cat. No. CC-3151) supplemented with 2 mM L-glutamine, 100 U/mL penicillin and streptomycin, 20 ng/mL recombinant human epidermal growth factor (Sigma Aldrich; Cat. No. E9644), 10 ng/mL recombinant human basic fibroblast growth factor (R&D Systems; 233-FB-025) and 1× B27 supplement (Invitrogen; Cat. No. 17504-044) (51). Cells were plated at a density of 20,000 cells or 30,000 cells per well for the MDA-MB-231 and Cal51 cell lines respectively in a 6-well low attachment plate (Corning; Cat. No. CLS3814) with 3mL of media. Fresh media were added to the wells every three days over the course of the assay. Mammospheres were measured four weeks post plating for the MDA-MB-231 cell lines and two weeks post plating for Cal51 cell lines.

### Drug sensitivity assays

To quantify a cell line’s sensitivity to a particular drug, 10,000 A375, DLD1, or MDA-MB-231 cells or 5,000 Cal51 cells were plated in 100μL of media in an 8 × 4 matrix on a flat-bottomed 96-well plate (Corning; Cat. No. 3596). Cells were allowed 24 hours to attach, then fresh media was added to each well. The highest concentration of a drug was added onto the first row of cells and then six three-fold serial dilutions were performed. Cells were grown in the presence of the drug for 72 hours then trypsinized and counted using a MacsQuant Analyzer 10 (Milltenyi Biotec). HTH-01-091 was a kind gift of Hubert Huang and Nathanael Gray (Dana-Farber Institute). Idarubicin, Oxaliplatin, Paciltacxel and Bleomycin were obtained from Selleck Chemcials (Cat No. S1228, S1224, S1150, and S1214). Fluorouracil was obtained from Sigma Aldrich (Cat. No. F6627-1G).

### Western blot analysis

Whole cell lysates were harvested using RIPA buffer (25 mM Tris, pH 7.4, 150 mM NaCl, 1% Triton × 100, 0.5% sodium deoxycholate, 0.1% sodium dodecyl sulfate, protease inhibitor cocktail, and phosphatase inhibitor cocktail). Protein concentration was quantified using the RC DC Protein Assay (Bio-Rad; Cat. No. 500-0119) or the Pierce BCA Protein Assay Kit (Thermo Fisher Scientific; Cat. No. 23225). Equal amounts of lysate were denatured and loaded onto an 8% SDS-PAGE gel. The protein was transferred onto a polyvinylidene difluoride membrane using the Trans-Blot Turbo Transfer System (Bio-Rad). For westerns with phospho-antibodies, membranes were blocked in 5% BSA, while all other antibodies were blocked with 5% milk. The following antibodies and dilutions were used: Anti-MELK (Abcam; Cat. No. ab108529) at a dilution of 1:3000, Anti-elF4B (Cell Signal; 3592) at a dilution of 1:2000, Anti-ASK1 (Abcam, Cat. No. ab45178) at a dilution of 1:1000, Anti-cyclin B (Abcam; Cat. No. ab32053) at a dilution of 1:10000, Anti-Phospho-ASK1 (Cell Signal; 3765) at a dilution of 1:1000, Anti-Phospho-elF4B (Cell Signal; Cat. No. 5399) at a dilution of 1:1000, or anti-MCL1 (Cell Signal; Cat. No. 5453) at a dilution of 1:2000. Blots were incubated with the primary antibody overnight at 4°C. Anti-alpha tubulin (Sigma-Aldrich; Cat. No. T6199) at a dilution of 1:20000 or anti-GAPDH (Santa Cruz Biotechnology; Cat. No. sc-365062) at a dilution of 1:20000 were used as loading controls. Membranes were washed at room temperature for an hour before they were incubated in secondary antibodies [Anti-Rabbit (Abcam; Cat. No. ab6721) at 1:50000 for anti-MELK and at 1:20000 for all other antibodies or Anti-mouse (Bio-Rad; Cat. No. 1706516) at 1:50000 for anti-tubulin and anti-GAPDH)] for an hour.

### Xenograft growth assays

Nude mice were obtained from The Jackson Laboratory (Cat. No. 002019). To perform the xenograft injections, Cal51 and MDA-MB-231 MELK-KO and Rosa26 clones were harvested and resuspended at a concentration of 10^7^ cells/mL in 1× cold PBS. The cell suspension was then mixed 1:1 with growth factor reduced-matrigel (Corning; Cat. No. 47743-720). Each mouse was injected subcutaneously in the left and right flanks with 100μL of the cell suspension, containing 5×10^6^ cells. Tumors were monitored by visual inspection routinely after injection. Once a tumor was visible, mice were measured every three days by caliper in duplicate. Tumor volume was calculated using the formula V = 1/2 (longer axis)(shorter axis)^2^. All mouse protocols were approved by the CSHL Institutional Animal Care and Use Committee.

### CRISPR plasmid construction and virus generation

Guide RNAs were previously described in Lin et al. (15). In short, oligonucleotides were cloned into the LRG 2.1 vector [a gift from Jun-Wei Shi (University of Pennsylvania) and Chris Vakoc (Cold Spring Harbor Laboratory)] using a BsmBI digestion (52). To produce virus, HEK293T cells were transfected using the calcium-phosphate method. Supernatant was harvested 48 to 72 hours post-transfection, filtered through a 0.45μm syringe, and then applied to cells with 4μg/mL polybrene (49). Guide RNAs in each clone are listed in Table S1.

### Analysis of CRISPR-mediated mutagenesis

Single cells isolated via fluorescence-activated cell sorting (FACS) were grown into clonal populations. Genomic DNA was extracted from these populations with the QIAmp DNA Mini kit (Qiagen; Cat. No. 51304). The cut site regions targeted by the guide RNAs were amplified using the primers listed in Table S2. PCR products were then sequenced with the forward and reverse primer at the Cold Spring Harbor Laboratory sequencing facility.

### Analysis of published gene expression data

Data from normal human tissue was acquired from (53). Data from senescent fibroblasts was acquired from (54). Data from stimulated lymphocytes was acquired from (55). Cancer patient cohorts and probeset definitions were downloaded from the Gene Expression Omnibus as described in Table S3 (56). Probesets annotated to the same gene were collapsed by averaging. Data were cleaned and processed using python’s pandas library, and cox proportional hazard models were constructed using the survival library in R and as described in (17). Several additional python packages were used for clarity, conciseness, and correctness including numpy, getopt and rpy2. Source code is available on github (https://github.com/joan-smith/survival-analysis-scripts).

---

**Supplemental Figure 1.**
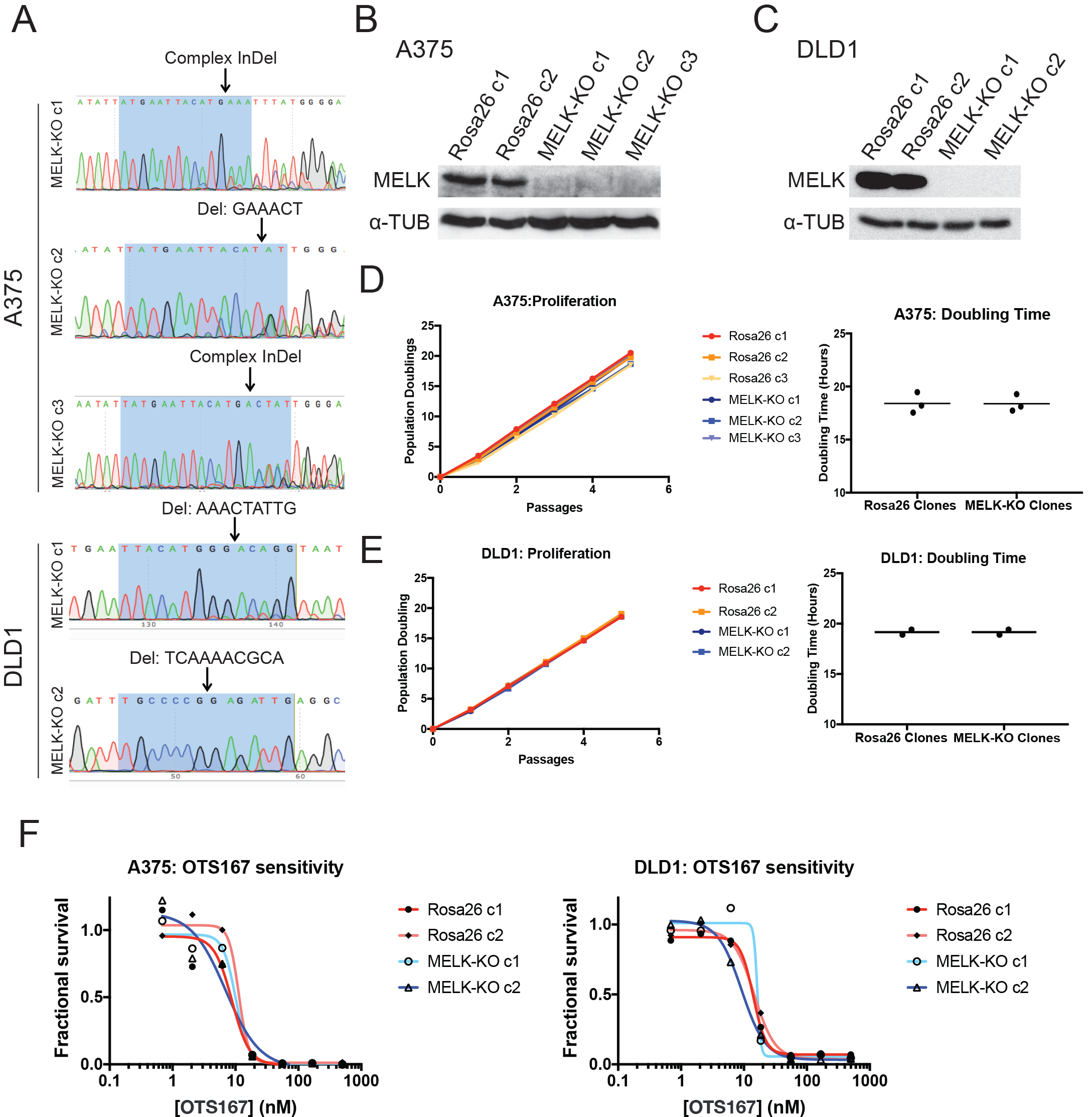
Generation and characterization of MELK-KO clones in A375 and DLD1. (A) Sanger sequencing of MELK knockout clones in the DLD1 (colorectal cancer) and A375 (melanoma) cell lines. Highlighted regions indicate bases recognized by the gRNA. (B-C) Western blot analysis of DLD1 and A375 MELK-KO clones. (D-E) Proliferation and doubling time analysis of DLD1 and A375 control and MELK-KO clones. (F) Dose-response curves of the putative MELK inhibitor OTS167 in A375 and DLD1 control and MELK knockout clones.

**Supplemental Figure 2.**
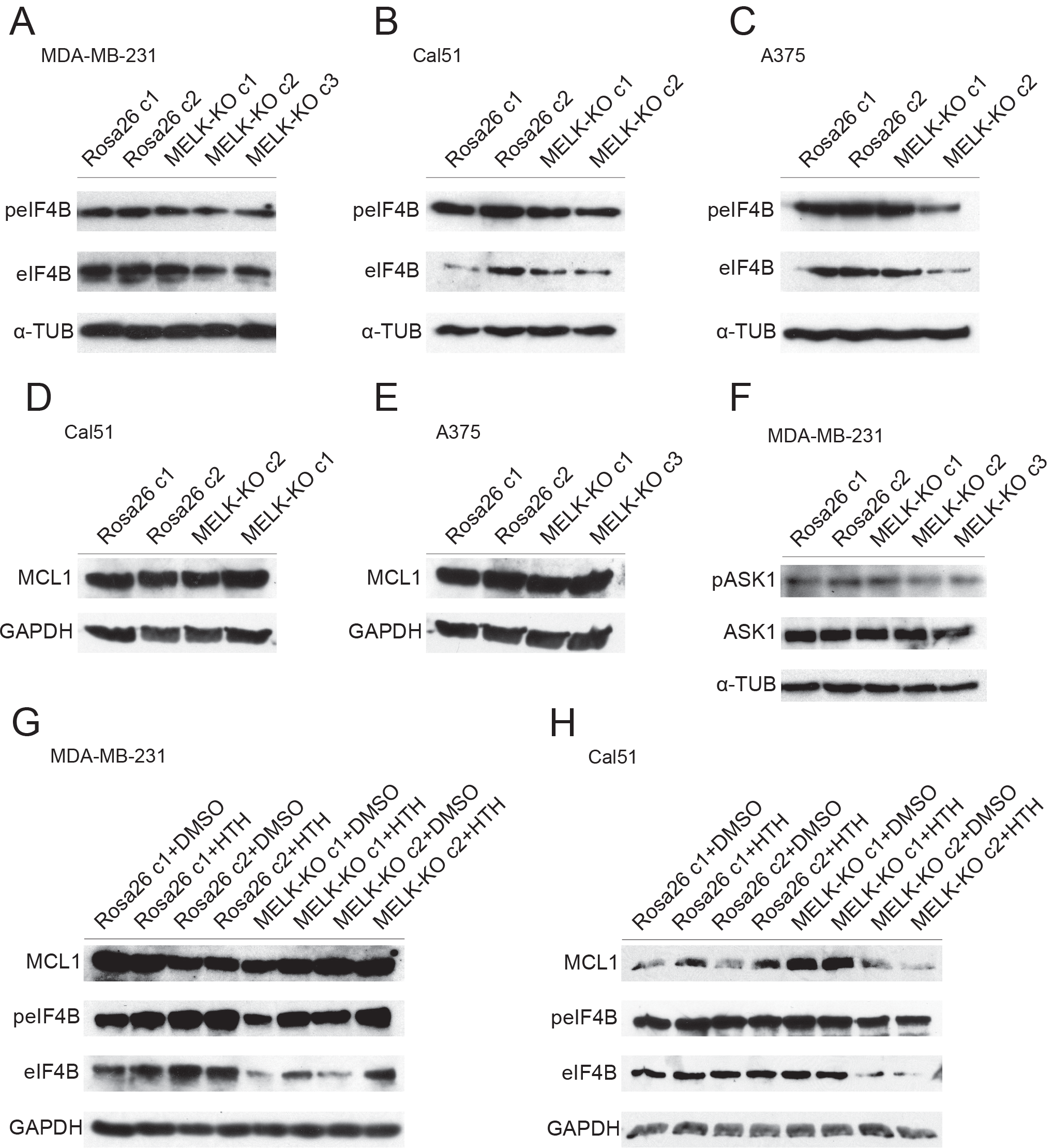
MELK is not required for the phosphorylation or expression of previously-reported targets. (A-C) Western blot analysis of MDA-MB-231, Cal51, or A375 with antibodies against phospho-EIF4B and total EIF4B. (D-E) Western blot analysis of Cal51 or A375 with antibodies against MCL1. (F) Western blot analysis of MDA-MB-231 with antibodies against phospho-ASK1 and total ASK1. (G-H) Western blot analysis of MDA-MD-231 or Cal51 treated with 1μM of HTH-01-091 with antibodies against MCL1, phospho-EIF4B and total EIF4B.

